# A NEW POSTURAL MOTOR RESPONSE TO SPINAL CORD STIMULATION: POST-STIMULATION REBOUND EXTENSION

**DOI:** 10.1101/2024.06.13.598885

**Authors:** Amr A. Mahrous, Matthieu Chardon, Michael Johnson, Jack Miller, CJ Heckman

**Affiliations:** Department of Neuroscience, Feinberg School of Medicine, Northwestern University, Chicago, IL, USA; Department of Physical Therapy and Human Movement Sciences, Feinberg School of Medicine, Northwestern University, Chicago, IL, USA; Department of Physical Medicine and Rehabilitation, Feinberg School of Medicine, Northwestern University, Chicago, IL, USA; Shirley Ryan AbilityLab, Chicago, IL, USA

**Keywords:** Spinal cord stimulation, Rebound excitation, Limb extension, Sit-to-stand transition, Rehabilitation

## Abstract

Spinal cord stimulation (SCS) has emerged as a therapeutic tool for improving motor function following spinal cord injury. While many studies focus on restoring locomotion, little attention is paid to enabling standing which is a prerequisite of walking. In this study, we fully characterize a new type of response to SCS, a long extension activated post-stimulation (LEAP). LEAP is primarily directed to ankle extensors and hence has great clinical potential to assist postural movements. To characterize this new response, we used the decerebrate cat model to avoid the suppressive effects of anesthesia, and combined EMG and force measurement in the hindlimb with intracellular recordings in the lumbar spinal cord. Stimulation was delivered as five-second trains via bipolar electrodes placed on the cord surface, and multiple combinations of stimulation locations (L4 to S2), amplitudes (50-600 uA), and frequencies (10-40 Hz) were tested. While the optimum stimulation location and frequency differed slightly among animals, the stimulation amplitude was key for controlling LEAP duration and amplitude. To study the mechanism of LEAP, we performed in vivo intracellular recordings of motoneurons. In 70% of motoneurons, LEAP increased at hyperpolarized membrane potentials indicating a synaptic origin. Furthermore, spinal interneurons exhibited changes in firing during LEAP, confirming the circuit origin of this behavior. Finally, to identify the type of afferents involved in generating LEAP, we used shorter stimulation pulses (more selective for proprioceptive afferents), as well as peripheral stimulation of the sural nerve (cutaneous afferents). The data indicates that LEAP primarily relies on proprioceptive afferents and has major differences from pain or withdrawal reflexes mediated by cutaneous afferents. Our study has thus identified and characterized a novel postural motor response to SCS which has the potential to expand the applications of SCS for patients with motor disorders.

## INTRODUCTION

Animal studies, particularly in feline preparations, have demonstrated the remarkable capacity of the spinal cord to orchestrate functional motor activity independently from the brain. Pioneering experiments conducted by Sherrington and colleagues in the early 20^th^ century demonstrated the potential for electrical stimulation of the spinal cord to elicit locomotor movements in the absence of any descending inputs (Roaf and Sherrington, 1910). Concurrent investigations by Brown (1911) further underscored the intrinsic capabilities of the spinal cord to generate locomotor-like activity in spinalized, deafferented preparations. Therefore, spinal cord stimulation (SCS) was used decades later to evoke locomotion following acute spinal cord injury (SCI) in anesthetized cats (Iwahara et al., 1991).

These seminal studies laid the groundwork for the use of SCS to rehabilitate motor functions in patients with SCI. Indeed, non-weight-bearing locomotor-like activity can be evoked in humans paralyzed for years by SCI while in a supine position (Dimitrijevic et al., 1998; Minassian et al., 2004). More recently, the integration of epidural SCS along with rehabilitative training showed potential in restoring weight-bearing stepping in patients with SCI (Harkema et al., 2011; Wagner et al., 2018; Lorach et al., 2023)

Despite these remarkable achievements in leveraging SCS to promote locomotor recovery, there remains a notable scarcity of studies that specifically investigate SCS-induced postural movements crucial for sit-to-stand transitions and standing stability. The restoration of stable posture represents a primary aspiration for individuals with SCI (Snoek et al., 2004), and offers several additional physiological benefits. Not only does it serve as a prerequisite for walking, but standing can also improve autonomic output, including cardiovascular, bowel, bladder, and sexual functions (Walter et al., 1999; Agarwal et al., 2003; Harkema et al., 2008; Biering-Sorensen et al., 2009).

While weight-bearing hindlimb extension elicited by intraspinal microstimulation has been previously reported in feline models (Mushahwar et al., 2000; Tai et al., 2003) and postural limb reflexes have been shown to be facilitated by epidural stimulation in spinalized rabbits (Musienko et al., 2010), little attention has been directed towards evoking analogous postural responses in humans with SCI. Noteworthy exceptions include the induction of non-weight-bearing leg extension in SCI patients using low-frequency epidural SCS (Jilge et al., 2004b) and the facilitation of self-assisted standing via low-frequency transcutaneous SCS (Sayenko et al., 2019).

In the current study, we report, in a feline preparation, a new response to SCS selectively directed to ankle extensor muscles which are crucial for postural stability (Hirono et al., 2020). This response is robust, reproducible, and controllable via stimulation amplitude and hence can be harnessed clinically to facilitate sit-to-stand transition and other postural movements. It manifests as a sustained extension which starts immediately after a short stimulation train; hence we refer to it as long extension activated poststimulation (LEAP). Our single-neuron intracellular recordings indicate that this response is mediated by a network of interneurons and involves activation of multiple types of afferents. To fully characterize this response, we have examined combinations of stimulation frequencies, amplitudes, and locations along the lumbosacral cord. The isometric forces measured from the ankle extensors during LEAP are high and can support full weight bearing in cats. Hence, this new response can be adopted clinically to facilitate postural movements in patients with motor disabilities.

## RESULTS

To examine spinal cord stimulation (SCS) protocols that can support postural movements, we employed an *in vivo* setup utilizing decerebrate cats (Fig. 1A). This animal model enabled us to study the responses to SCS without the confounding effects of anesthetics. Bipolar electrodes were precisely positioned on the left side of the cord dorsum targeting multiple locations marked by dorsal root entry zones spanning from L4 to S2 (Fig. 1B). Electrical stimulation was delivered as trains of 1 ms pulses at multiple frequencies (10, 20, and 40 Hz) and amplitudes (1-10 times the motor threshold, Fig. 1C). Motor responses were monitored via EMG and force transducers, focusing on the ankle flexor tibialis anterior (TA), and the ankle extensor soleus (SOL) ipsilateral to stimulation.

**Fig. 1:**
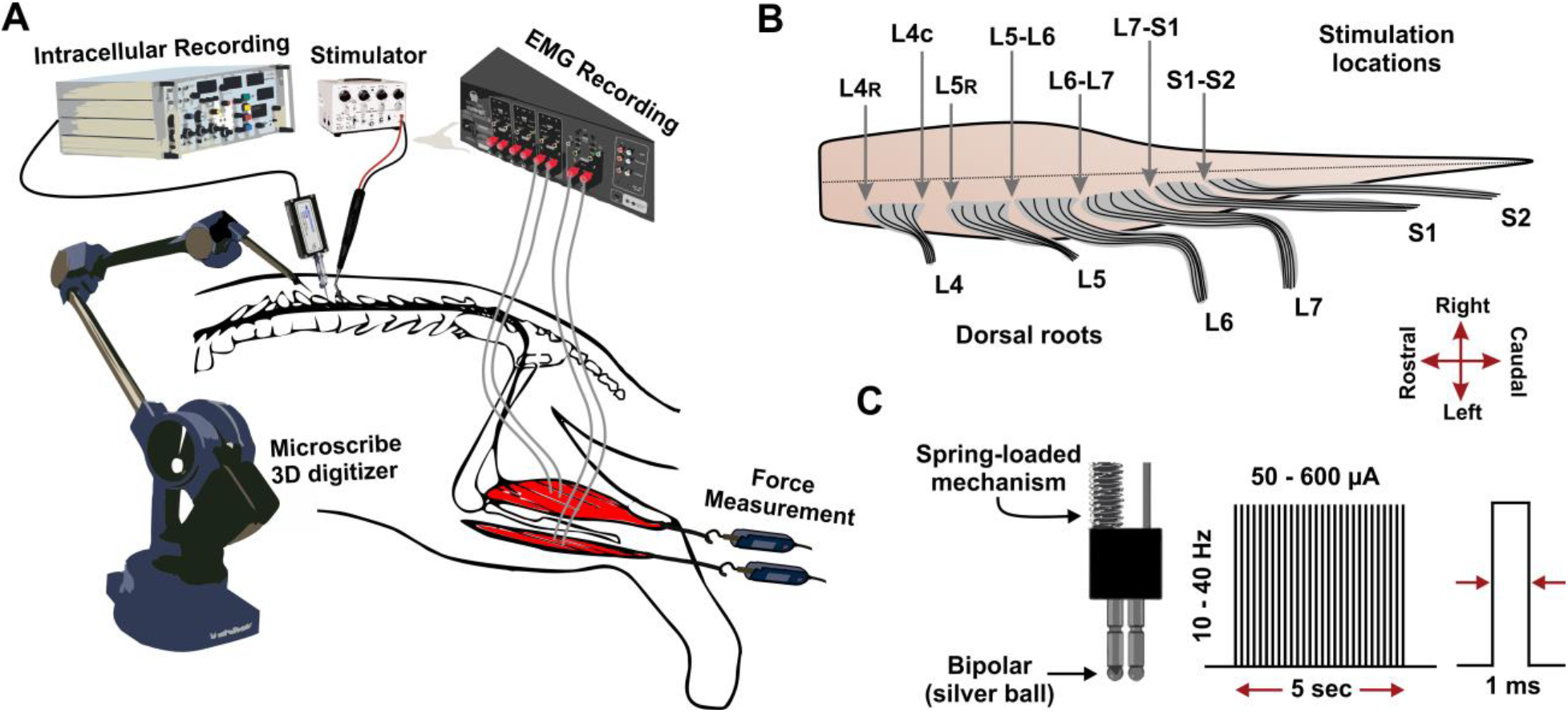
Experimental setup for spinal cord stimulation. **A:** Schematic of the *in vivo* preparation setup in decerebrate cats. The response to lumbar spinal cord stimulation was recorded in ankle flexors and extensors (isometric force and EMG) and intracellularly in spinal motoneurons and interneurons. Each stimulation location was digitized using a Microscribe MLX Digitizer system and recorded alongside electrophysiology data. **B:** Diagram of the lower lumbar and upper sacral segments of the cat spinal cord showing different stimulation locations. The two leads of the stimulation electrode (1 mm apart) were placed at junctions between dorsal root entry zones, such as L5-L6, on the left side of the cord. When the roots were distinctly separated by a distance, the two ends were designated as separate locations, such as the caudal end of L4 (L4C) and rostral end of L5 (L5R). **C:** Spinal cord stimulation electrode design and stimulation protocol. The bipolar silver ball electrode was loaded on springs that help maintain contact with the tissue while minimizing tissue damage due to any vertical movements. Stimulation trains were delivered for 5 seconds at 10, 20 or 40 Hz and with an amplitude of 50 – 600 µA (with 50 - 100 µA increments).

Certain combinations of stimulation parameters evoked a distinctive pattern of muscle activation around the ankle, characterized by robust persistent contraction of the extensors after the stimulation train ended (Fig. 2A). This post-stimulation extensor response is the focus of this study, as it has potential clinical benefit in supporting sit-to-stand transition and other postural movements.

**Fig. 2:**
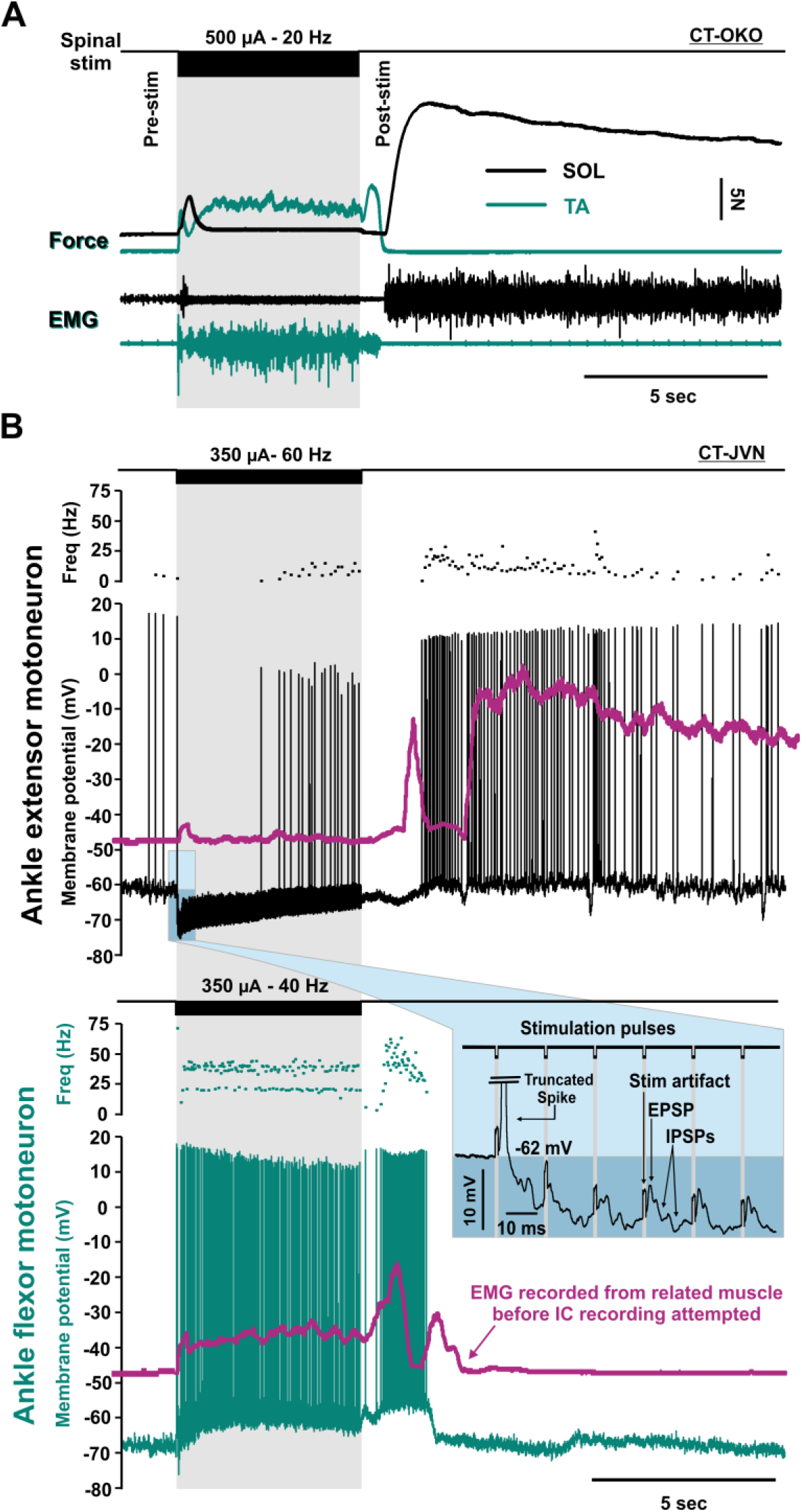
Long extension activated poststimulation (LEAP) from the cat spinal cord. **A:** Force and EMG recorded from soleus (SOL, black) and tibialis anterior (TA, green) of the left hindlimb in a decerebrate cat in response to a single five-second train (shaded area) of subdural electrical stimulation at the junction between L6 and L7 dorsal root entry zones. Note that stimulation is delivered to the left side of the cord ipsilateral to the recorded muscles. Soleus muscle responds with a strong and prolonged contraction shortly after stimulation train ends. N= 19 experiments. **B:** Intracellular recordings from a different experiment showing typical firing behavior of an ankle extensor motoneuron (black, n= 55 triceps MNs) and ankle flexor motoneuron (green, n= 14 common peroneal MNs) during and after stimulation. Each of the two cells were recorded several hours apart in the same preparation. The superimposed traces (purple) are the integrated EMG activity of soleus and TA recorded few hours before paralysis was induced for intracellular recording stability. ***Inset:*** A close-up on motoneuron membrane potential at the beginning of the stimulation train. Some triceps motoneurons fire an action potential following the first pulse of stimulation followed by hyperpolarization during most of the train time.

### Long extension activated poststimulation (LEAP)

During brief stimulation trains (5 seconds throughout this study), the flexor muscle TA was activated throughout the train with occasional brief post-stimulation contraction (fig. 2A). Conversely, the ankle extensor SOL was not active during stimulation except for a brief twitch at the beginning of the train. However, after stimulation ended, the ankle extensors generated a robust sustained contraction lasting several seconds (fig. 2A). This extensor response was ipsilateral to stimulation and primarily restricted to the ankle, and therefore distinct from a crossed extension reflex.

The same phenomenon recorded intracellularly from both ankle extensor and flexor motoneurons in another animal is shown in Fig. 2B. We recorded a total of 55 tibial nerve motoneurons (extensor MNs), 19 common peroneal motoneurons (flexor MNs), and 11 un-identified motoneurons. Approximately 73% of the impaled flexor motoneurons fired repetitively and their membrane potential remained depolarized throughout the stimulation train (n= 14/19 MNs, Fig. 2B). Conversely, 71% of the impaled extensor motoneurons exhibited hyperpolarization during stimulation (fig. 2B, n= 39/55 MNs). Following each pulse of stimulation, an EPSP was observed followed by larger and longer IPSPs which resulted in an overall hyperpolarization of the membrane (fig. 2B, inset). The EPSP elicited by the first pulse of the train preceded any hyperpolarization and thus frequently resulted in an action potential (n= 25/55 cells), accounting for the brief twitch observed in SOL at the beginning of stimulation (Fig 2A). Shortly after stimulation ended, 89% of the extensor motoneurons recorded (n= 49/55) started to depolarize/fire for several seconds.

Here, we term this post-stimulation activity in extensor motoneurons/muscles as long extension activated poststimulation (LEAP). This LEAP response was relatively consistent across experiments and reproducible for several hours within each experiment. Throughout this study, SCS was delivered directly to the spinal cord surface after opening the dura (subdural). However, the LEAP response can also be evoked by epidural stimulation (Suppl. Fig. 1).

### Stimulation parameters to evoke LEAP

We investigated the specific stimulation parameters needed to evoke LEAP in the ankle extensors, including stimulation location, frequency, and amplitude. The dorsal root entry zones served as reliable landmarks for precise and consistent placement of the stimulation electrode across animals (See Fig. 1B). These locations were carefully digitized in each experiment by tracing the dorsal root entry zones, the border of laminectomy, and the position of the electrode tip using a Microscribe MLX 3-D Digitizer system. An example of such a data set is shown in Suppl. Fig. 2.

Anatomical studies have located the motor pools of the cat TA and SOL at L6 and L7 segments (Yakovenko et al., 2002). Hence, we stimulated multiple sites spanning from L4 to S2 segments in 10 animals (Fig. 3). While LEAP responses could be evoked at multiple stimulation locations (Fig. 3A), there was a noticeable variation in the response amplitude. The most pronounced LEAP responses were consistently observed either at L5-L6 segments (N= 5) or more caudally at L7- S1 (N=5, Fig. 3B). Notably, larger LEAP responses were observed at stimulation locations that had large TA responses and small soleus responses during the stimulation train (Suppl. Fig. 3).

**Fig. 3:**
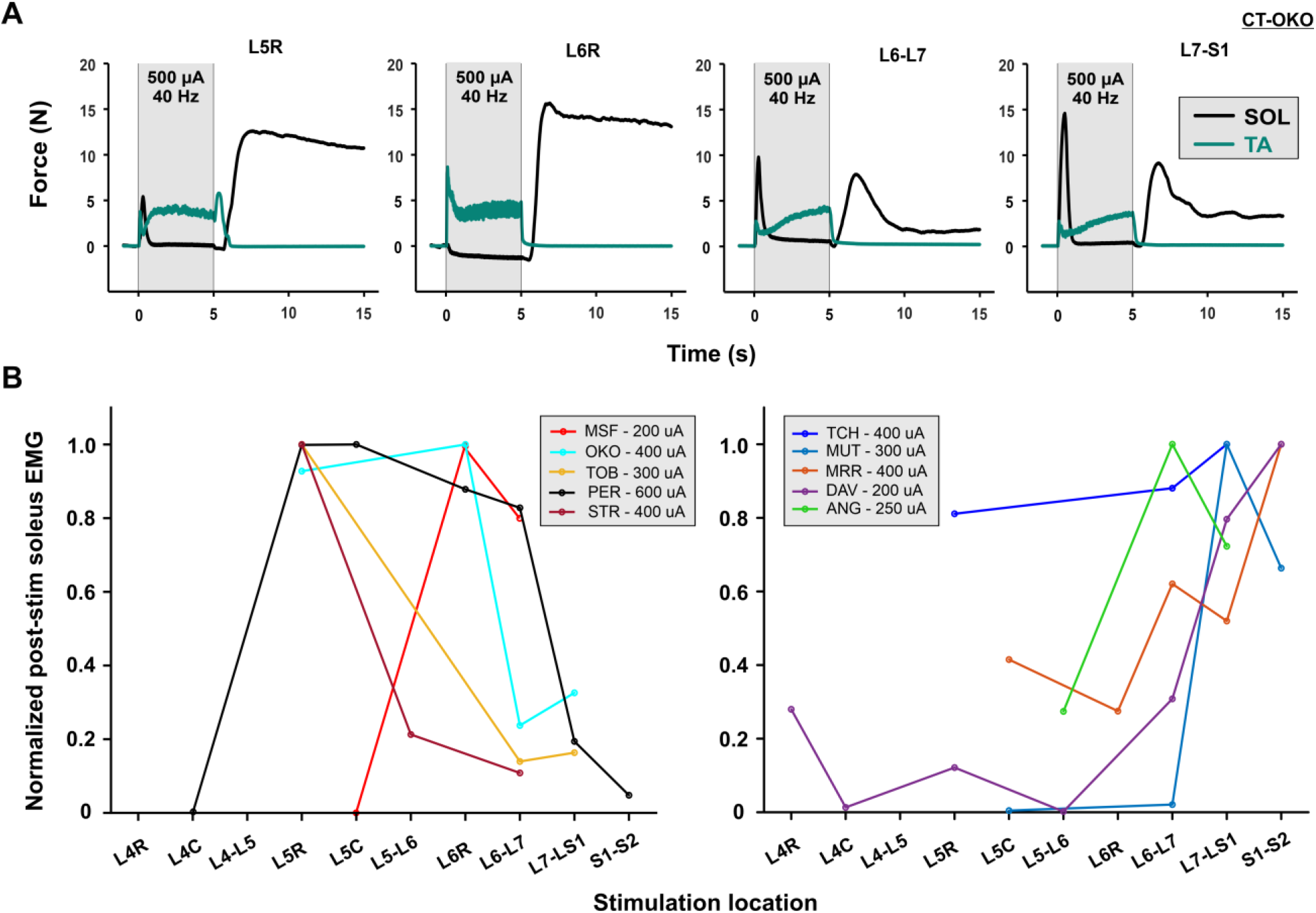
Optimum stimulation location for evoking LEAP. **A:** Examples of force measurements from SOL and TA during and after stimulation at 4 different locations in the lumbosacral area in one experiment. The amplitude of the LEAP response varies noticeably at different locations. **B:** Summary of the effect of stimulation locations on LEAP amplitude in 10 experiments (different colors). The response amplitude was measured as the integrated EMG response in soleus in the 30 seconds following the stimulation train and normalized to the maximum response evoked in each experiment. A near maximum stimulation amplitude at 40 Hz was used. The optimum location (maximum response amplitude) varied slightly among animals. The data was divided into 2 groups with the optimum location around the lumbosacral junction (right panel) or strictly in the lumbar segments (left panel).

Additionally, we tested multiple stimulation frequencies including 10, 20, and 40 Hz while maintaining a fixed 5-second stimulation duration. Submaximal stimulation amplitudes were selected during these trials to ensure that spinal circuits are not saturated with current at higher frequencies. In 5 out of 8 experiments, the response increased linearly with increasing frequency (Fig. 4), while the other 3 experiments had the maximal response at 20 Hz. Generally, the data indicates that LEAP can be evoked at a broad range of stimulation frequencies and might be used to control LEAP response at submaximal stimulation amplitudes.

**Fig. 4:**
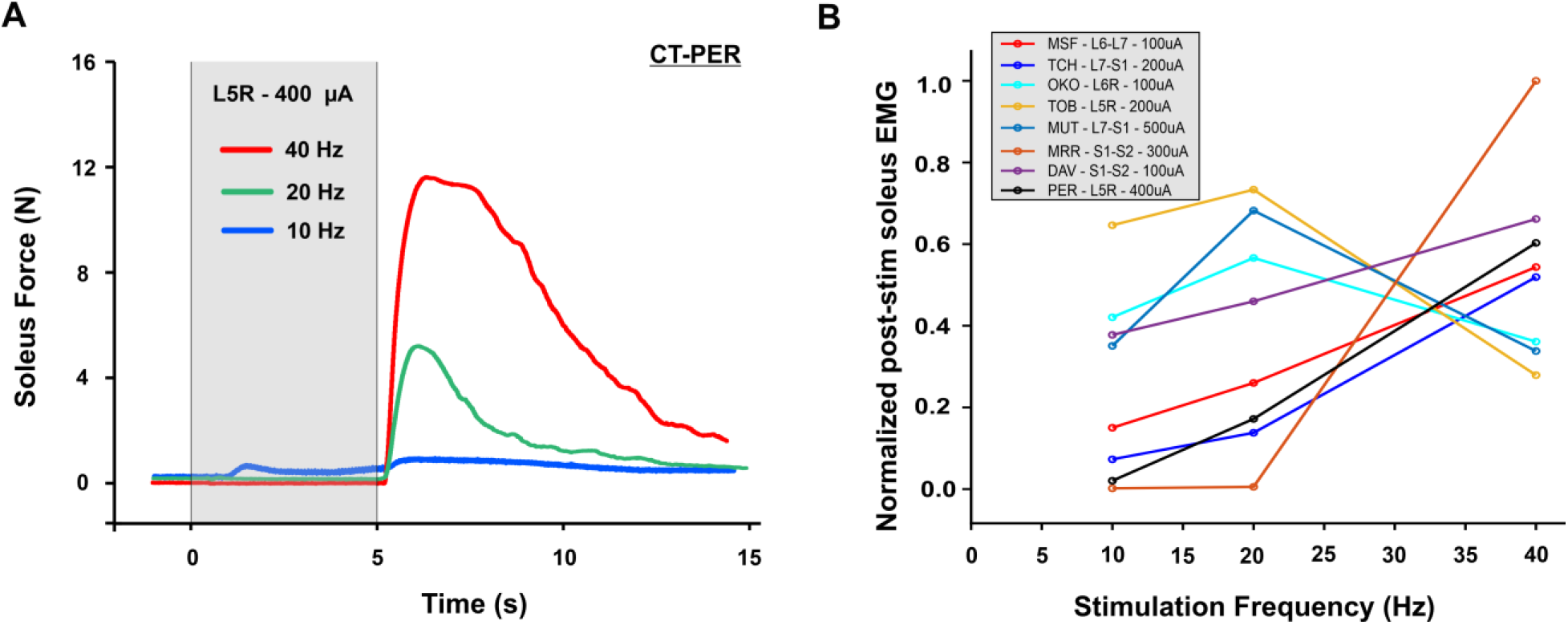
The effect of stimulation frequency on LEAP. **A:** Examples of force traces measured from SOL during and after stimulation at the optimum stimulation location for that animal (L5R) and stimulation amplitude that produces about half maximum response at 40 Hz (400 µA, see fig. 5). **B:** Summary of the effect of stimulation frequency (10, 20, and 40 Hz) on LEAP amplitude in 8 experiments (colors, same as in fig. 3). Electrical stimulation was delivered at an amplitude that produces about 50% of maximum response at 40 Hz in each experiment, and optimum stimulation location (chosen from fig. 3). The amplitude of LEAP was measured as the integrated EMG response in soleus in the 30 seconds following the stimulation train.

Stimulation amplitudes exceeding the motor threshold were generally required to evoke LEAP, with the motor threshold typically ranging between 50-100 µA. The maximum LEAP was elicited at 2-5 times higher than the motor threshold with possible gradation of response observed between these values (Fig. 5 and Suppl Fig. 4). Overall, the amplitude of LEAP response was readily controllable via stimulation amplitude modulation.

**Fig. 5:**
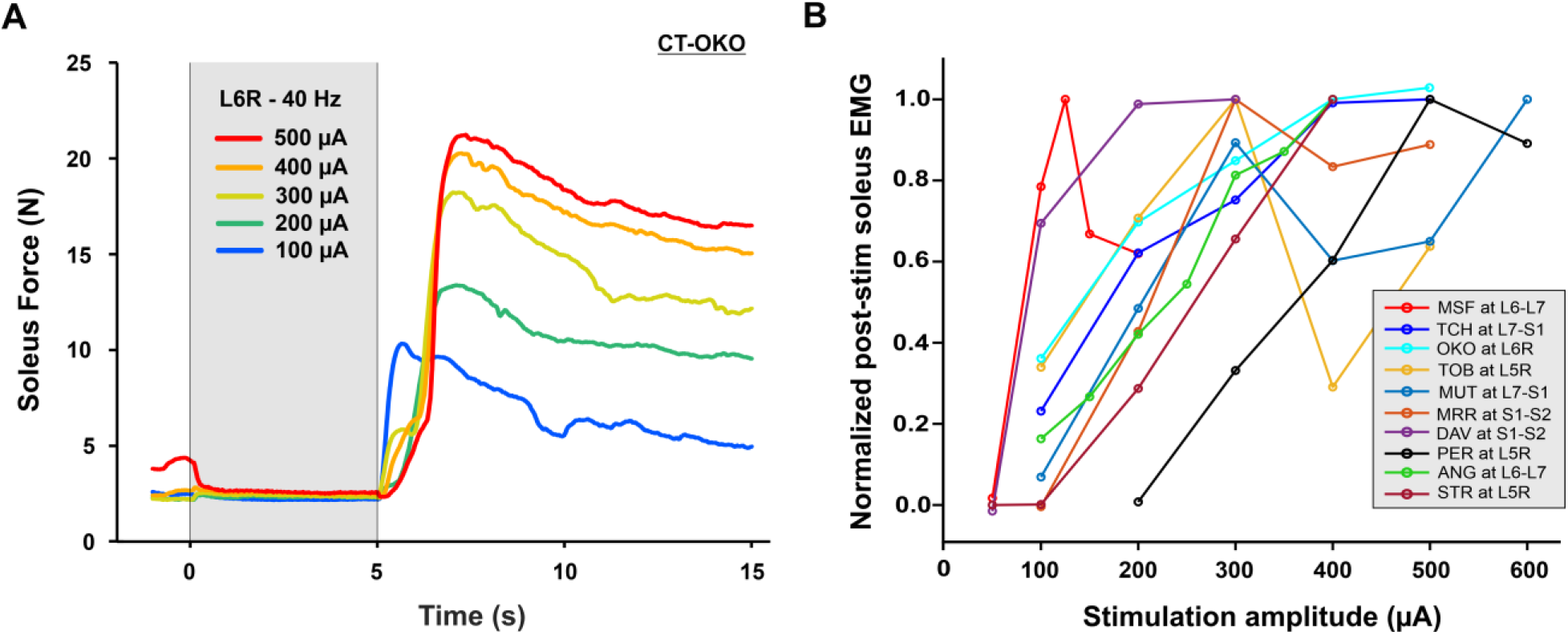
Control of LEAP using stimulation amplitude. **A:** Examples of SOL force measurements during and after stimulation at the optimum stimulation location for that animal (L6R) and 40 Hz stimulation frequency. The amplitude of the LEAP response can be titrated using stimulation amplitude. **B:** Summary of the effect of stimulation amplitude (50-600 µA) on LEAP amplitude in 10 experiments (colors, same as in fig. 3 and 4). Electrical stimulation was delivered at 40 Hz to the optimum stimulation location in each experiment (chosen from fig. 3). LEAP amplitude was measured as the integrated EMG response in soleus in the 30 seconds following the stimulation train. The LEAP response can be controlled using the stimulation amplitude.

Therefore, LEAP response can be quickly established by scanning a few stimulation locations within the lower lumbar segments at an amplitude of 2-5xT and 40 Hz, followed by creating a dose-response relation at the optimal stimulation location.

### Is LEAP intrinsic to motoneurons or mediated by synaptic currents?

The prolonged post-stimulation firing in extensor motoneurons during LEAP could be caused by an excitatory synaptic current and/or an intrinsic self-sustained motoneuron firing mediated by ion channels that conduct persistent inward currents (PICs) (Heckman et al., 2005). To investigate the mechanism, we performed intracellular recordings of triceps motoneurons and repeatedly evoked LEAP while varying the membrane potential of the impaled cells. The peak depolarization during LEAP was then measured for each holding potential. At more hyperpolarized potentials, most triceps motoneurons showed increased depolarization during LEAP (fig. 6A). In 7/10 MNs, there was a negative correlation between the holding potential and the peak depolarization during

**Fig. 6:**
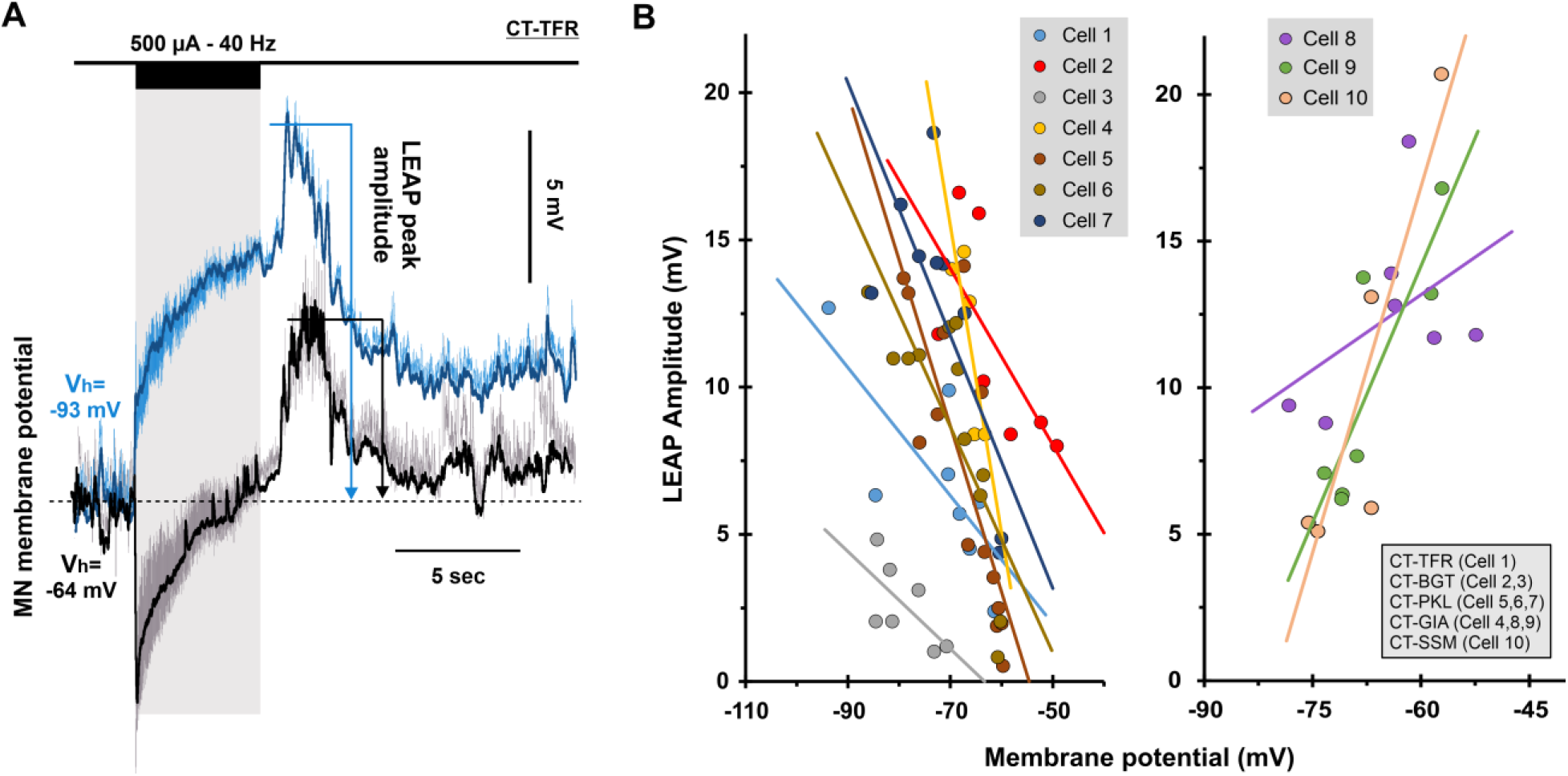
Motoneuron firing during LEAP response is driven by a combination of synaptic excitation and intrinsic mechanisms. **A:** Intracellular recording of LEAP from a triceps (TS) motoneuron at resting potential (black) and when hyperpolarized by a DC current (blue). The traces were low-pass filtered to eliminate spikes and stimulation artifacts. LEAP was not inhibited by hyperpolarization of the membrane, but instead became larger. **B:** Summary of the peak amplitude of LEAP potential at different holding potentials measured in 10 TS motoneurons recorded in 5 experiments (legend). **Left**: 7/10 MNs exhibited negative correlation between LEAP and holding potential indicating that LEAP was mediated by an excitatory synaptic input in these cells. **Right**: in 3/10 MNs, the correlation was positive indicating that this group intrinsically-sustained their firing during the post-stimulation period.

LEAP (Fig. 6B). This indicates that LEAP is largely mediated by an excitatory synaptic current. Nonetheless, in 3/10 cells, there was a positive correlation indicating that LEAP in these cells was driven by a voltage-gated membrane conductance. This suggests that during LEAP, extensor motoneurons are driven by excitatory synaptic currents that can recruit intrinsic PICs.

If LEAP is indeed driven by synaptic currents, it is expected that a subset of interneurons would exhibit altered firing patterns poststimulation. Indeed, intracellular recordings of spinal interneurons revealed an increase in firing activity concurrent with LEAP in about 27% of recorded cells (fig. 7A, n= 7/26 cells). Conversely, 46% of cells showed no change in firing (n= 12/26 cells), while a decrease in firing after stimulation occurred in 27% of cells (n= 7/26 cells).

**Fig. 7:**
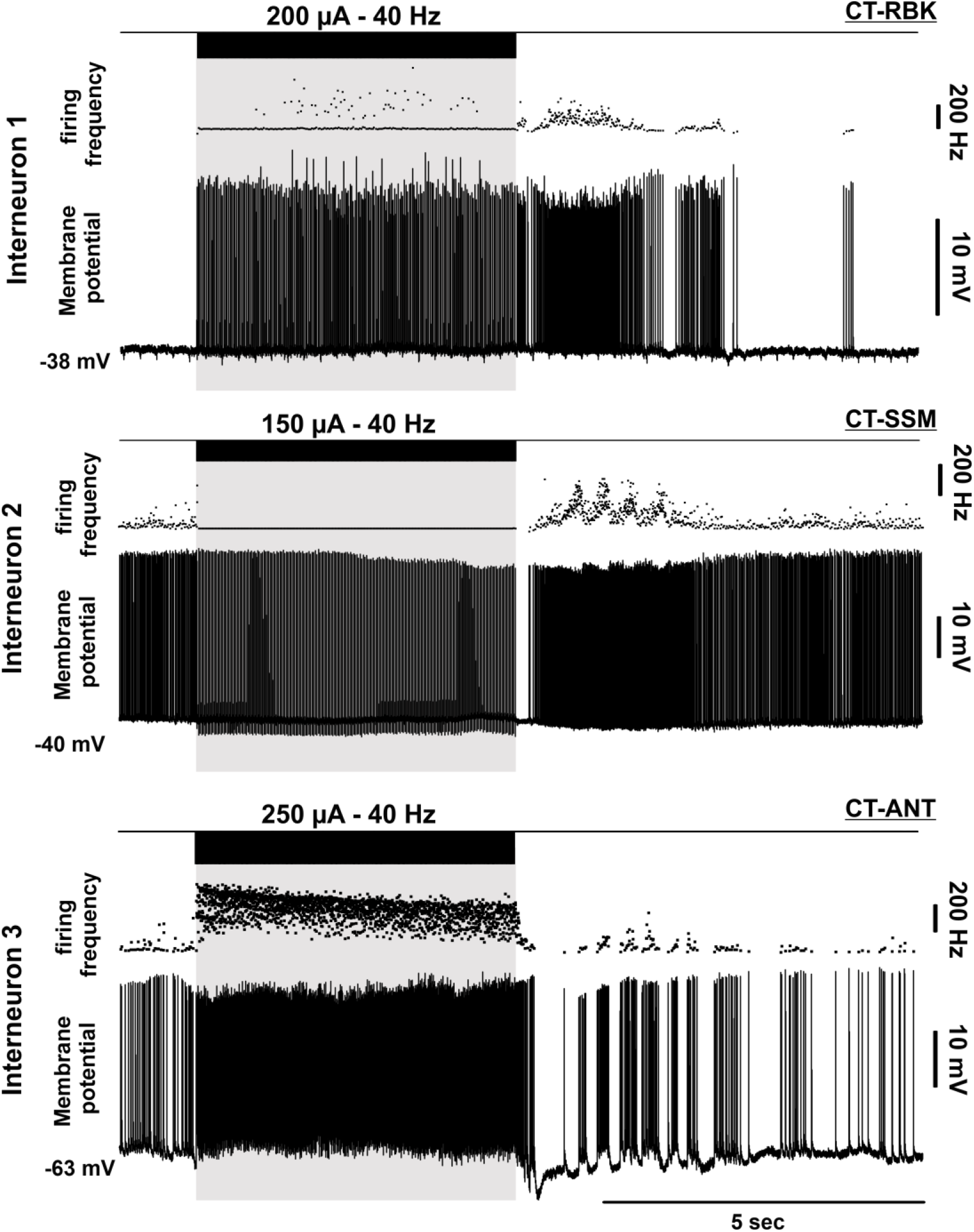
LEAP is mediated by spinal circuit activity. Intracellular recordings of spinal interneurons during LEAP using sharp glass microelectrodes. Even though the cells show distinct firing behaviors before and during spinal stimulation, they show increased firing in the post-stimulation period that coincide with LEAP (n= 7/26 INs, N=7 experiments).

### Types of sensory afferents involved in LEAP

In our stimulation protocol, the pulse width (1 ms) could activate both cutaneous and proprioceptive afferent axons (Szlavik and de Bruin, 1999). To discern the contribution of these sensory pathways to the activation of LEAP circuits, we modified the protocol to include shorter stimulation pulses (0.1 ms), which selectively recruit the largest 5% of fibers, primarily proprioceptive axons (Szlavik and de Bruin, 1999). Despite requiring higher stimulation amplitudes, the shorter pulse protocol successfully elicited LEAP responses (Fig. 8A), indicating the involvement of proprioceptive afferents.

**Fig. 8:**
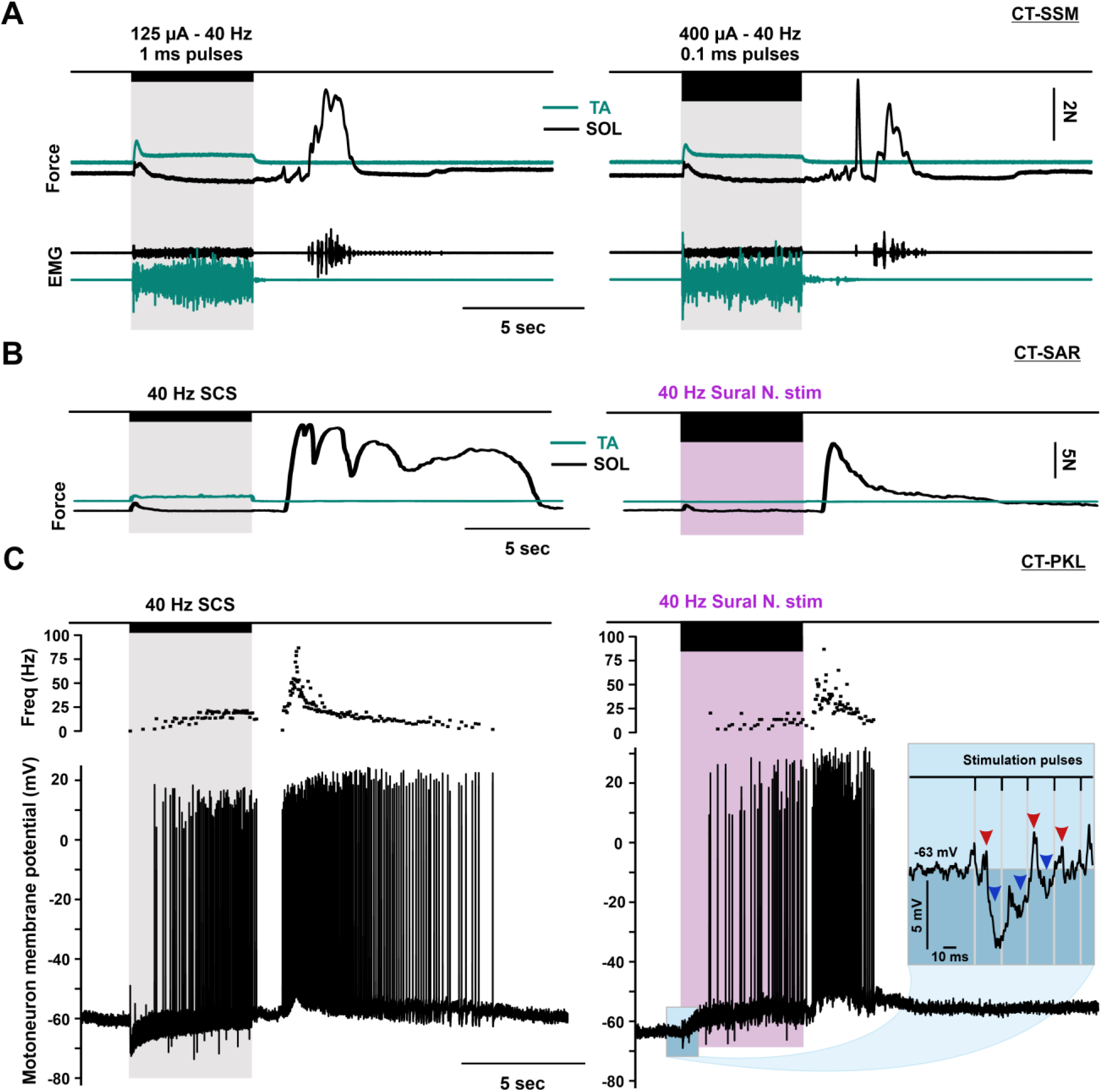
LEAP is mediated by multiple types of afferents. **A:** Force and EMG measurements from SOL and TA during and after stimulation at the same stimulation location and frequency but with different stimulation pulse width. Stimulation using short pulse width (0.1 ms, right), which is more selective for proprioceptive afferents, can evoke LEAP at higher stimulation amplitudes (N= 2 experiments). **B:** A smaller LEAP response can be evoked by peripheral stimulation of the ipsilateral sural nerve (mostly cutaneous fibers) using a bipolar cuff electrode. Same pulse width (1 ms) was used for central and peripheral stimulation (N= 3 experiments). **C:** Intracellular recordings of an ankle extensor motoneuron from a different experiment showing firing response of the same cell to stimulation of the spinal cord (left) and peripheral stimulation of the sural nerve (right). Inset shows the membrane response at the beginning of stimulation with mixed EPSPs (red arrowheads) and IPSPs (blue arrowheads).

To explore the role of cutaneous inputs, we stimulated an ipsilateral cutaneous nerve, the sural nerve, with same duration (5 sec), frequency (40 Hz), and pulse width (1 ms) using a nerve cuff. Sural stimulation required very high stimulation voltages (40-70 V) and generated smaller LEAP responses (Fig. 8B). When recorded intracellularly in extensor motoneurons (Fig. 8C, different experiment), sural stimulation evoked smaller IPSPs during the train compared to SCS, and the membrane potential was depolarized during most of the stimulation train which is distinctly different from SCS. Nevertheless, the data indicates that cutaneous afferents could contribute to LEAP. Therefore, LEAP circuit might be activated by multiple types of sensory afferents including proprioceptive and cutaneous pathways.

Collectively, our study has characterized a novel and robust extensor response elicited by spinal cord stimulation (SCS), delineated the specific stimulation protocol necessary for its induction, and elucidated its underlying mechanisms. With its reproducibility and reliability, this SCS-induced response holds promising clinical potential for assisting postural movements in individuals with motor disorders.

## DISCUSSION

SCS has become a major rehabilitation technique for many traumatic and degenerative conditions that affect central motor control, including spinal cord injury, stroke, spinal muscular atrophy, and Parkinson’s disease (Wagner et al., 2018; Milekovic et al., 2023; Powell et al., 2023; Prat-Ortega et al., 2024). In our study, we have identified a novel response to SCS, a long extension activated poststimulation (LEAP). The study of such post-stimulation responses not only offers insights into the underlying neural mechanisms of SCS but can also guide the optimization of stimulation parameters, thereby furnishing refined protocols to achieve specific motor outputs.

### The rebound nature of post-stimulation extension

The LEAP response seemed to follow a rebound excitation behavior, i.e. a depolarization that follows a period of hyperpolarization. This rebound behavior probably occurs in the interneurons that mediate this behavior and not directly in motoneurons. Nonetheless, triceps motoneurons that exhibited LEAP post-stim also showed noticeable hyperpolarization during stimulation. Similarly, EMG and force recordings show that this inhibition occurred throughout the motor pool, even turning off any background activity in the extensors that existed before stimulation indicating a widespread inhibition of extensors during the stimulation train. This is in line with our recent studies in the mouse spinal cord showing that repeated dorsal root stimulation recruits inhibitory pathways and plays a major role in short-term depression of the H-reflex (Mahrous et al., 2024).

It was also notable that the inhibition of extensors during stimulation was generally more pronounced at stimulation locations that elicited more flexor activation during stimulation and more LEAP post-stimulation (Suppl Fig. 3). Furthermore, evoked soleus activity was always opposite to that of the TA muscle. Taken together, this suggests that stimulation at these optimal locations probably recruits Ia afferents for the ankle flexors causing monosynaptic activation of the flexors and disynaptic reciprocal inhibition of their counterpart extensors (Feldman and Orlovsky, 1975). The post-stimulation activation of extensors, on the other hand, is probably mediated by post-inhibitory rebound of interneurons receiving similar Ia inhibition during stimulation, such as those belonging to the locomotor central pattern generators (Satterlie, 1985).

### Stimulation parameters for LEAP

The LEAP response was overall consistent and controllable through gradation of stimulation amplitude. However, stimulation effectiveness was location-dependent as it undoubtedly recruits sensory axons in the vicinity of the stimulation electrode (Capogrosso et al., 2013). Consequently, an optimal location for LEAP would be ideal for recruiting ankle flexor Ia afferents during stimulation and inducing post-inhibitory rebound that targets the extensors post-stimulation.

We have mapped the lower lumbar and upper sacral segments and found that the optimal stimulation location differs slightly among subjects but remains restricted to the lower lumbar segments (L5 to L7). These segments encompass the motor pools for the ankle flexors and extensors in the cat (Yakovenko et al., 2002), with larger flexor responses to stimulation of the more rostral part of this range and to surface stimulation than deeper stimulation with intraspinal electrodes (Tai et al., 2003; Tai et al., 2008). The variability of optimal location might stem from anatomical variability in the location of afferents. Similar variability was observed when mapping the cord dorsum potential while vibrating the triceps tendon in decerebrate cats (Mendez-Fernandez et al., 2019).

Human studies that focused on evoking lower limb extension found no location-specificity using epidural stimulation (Jilge et al., 2004b), but unexpectedly more specificity with transcutaneous stimulation (Sayenko et al., 2019). However, these human studies emphasized that low-frequency stimulation (5-15 Hz) is required for evoking extension while higher frequencies (20-30 Hz) evoke locomotor bursts (Jilge et al., 2004b; Sayenko et al., 2019). It is thought that low stimulation frequencies might preferentially activate extensors due to certain patterns of presynaptic inhibition and postsynaptic summation (Jilge et al., 2004a). Although we have not attempted to use frequencies higher than 60 Hz, we observed potent LEAP response across a broad range of frequencies (5-60 Hz). The possibility of evoking LEAP at high frequencies is advantageous because low frequencies can be uncomfortable for some patients (Sayenko et al., 2019). In addition, frequencies higher than 50 Hz can also help alleviate spasticity (Pinter et al., 2000).

### Sensory afferents that trigger LEAP

Short stimulation pulses that are more selective to proprioceptive afferents effectively triggered LEAP. The repeated activation of these afferents can evoke prolonged responses in spinal interneurons as has been shown with high-frequency tendon vibration, which specifically activate Ia afferents (Mendez-Fernandez et al., 2019). Furthermore, the clear inhibition of the extensors during the stimulation train is probably mediated by Ia inhibitory interneurons enabling alternating activity between the antagonistic muscle pair.

Even though high-intensity cutaneous nerve (sural) stimulation evoked some LEAP response, it was distinct from that evoked via SCS. This indicates that LEAP is not primarily mediated by cutaneous afferents. In addition, it is evident that LEAP lacks fundamental features of the withdrawal and pain reflexes mediated by cutaneous afferents. For example, LEAP involves ipsilateral extensor rather than flexor responses and lacks crossed extension response in the other limb (not shown). In addition, the flexor activity during the stimulation train was restricted to the ankle, with no knee flexor activity (Schouenborg and Kalliomaki, 1990). Finally, we observed no wind up of responses with repeated stimulation as it has been shown for C-fiber responses (Gozariu et al., 1997). Therefore, the data suggest that LEAP relies primarily on proprioceptive afferents with perhaps involvement of some cutaneous afferents but not pain pathways.

### Clinical significance

Postural stability in humans requires 5-20% of the maximum voluntary torque in the ankle (Hirono et al., 2020). Moreover, ankle extensors play a major role in gait stability during walking (Namayeshi et al., 2023). Our study has identified a new response to SCS that is selectively directed to ankle extensors. This response is controllable via modulation of stimulation amplitude and is directed mainly to ankle extensors; hence it has great potential in clinical contexts.

The force output from the SOL muscle during LEAP is potent, reaching up to 20 N at its peak, underscoring its utility in various motor tasks. In addition, it is coupled with synergistic effects from the gastrocnemius muscles. Although we have measured isometric forces and did not attempt to test if they can support weight bearing, we know that they exceed those of ground reaction forces of cat hindlimbs during standing and walking (Guillot et al., 2013; Corbee et al., 2014). Therefore, LEAP responses can generate significant ankle extension torque that can help with many motor tasks. However, it is imperative to acknowledge the limitations of this study, including the need for future investigations to validate LEAP under pathological conditions and the potential differences in muscle-specific activity between cats and humans. Nonetheless, LEAP presents a simple yet effective SCS-elicited response with multifaceted potential applications in augmenting plantar flexion and enhancing lower limb motor function.

## METHODS

### Animals

All experimental procedures were approved by the Northwestern University Institutional Animal Care and Use Committee. The data was obtained from 19 adult cats of either sex weighing 2.5-4 kg. All animals were obtained from a designated establishment for scientific research and were housed and fed within designated areas monitored daily by veterinary staff and trained personnel. Animals underwent acute terminal experiments, in which initial surgical procedures were done under deep gaseous anesthesia. A precollicular decerebration was then performed before data collection which allows the discontinuation of anesthesia (Silverman et al., 2005), thus relieving any suppressive effects on spinal circuits.

### Terminal surgical procedures

#### Anesthesia

Animals were initially anesthetized by inhalation of a mixture of isoflurane and N_2_O (4% in 100 % O_2_) in a custom chamber and then via a tracheal tube (1.5% – 3% in 100% O_2_). The right common carotid artery and right jugular vein were cannulated to monitor blood pressure, and deliver intravenous fluids, respectively. A heating lamp and circulating water pad were used to maintain body temperature which was continuously monitored using an esophageal thermometer. Throughout the surgery, body temperature, arterial blood pressure, heart rate, respiratory rates, reflexes, and muscle tone were checked/recorded every 15 minutes and used to adjust the level of anesthesia.

#### Hindlimb preparation

We dissected the left leg to expose the distal muscles and nerves around the ankle. A small incision was made in the front of the leg to expose the tibialis anterior (TA) muscle and separate it from surrounding tissue. The same procedure was done in the back of the leg to separate the medial gastrocnemius (MG), lateral gastrocnemius (LG), and soleus (SL). Surgical sutures were then tied securely around the TA and SOL distal tendons at the ankle, and the tendons were disarticulated to be connected later through sutures to force transducers. The skin incision was then extended at the popliteal fossa to expose the common peroneal (CP) and tibial nerves and a bipolar nerve cuff was placed around each for stimulation. Nerve stimulation was used to evoke antidromic action potentials in motoneurons to identify their target muscles during intracellular recordings. In some experiments, a cuff electrode was also placed around the sural nerve to test the effect of stimulating the cutaneous afferents. In addition, we inserted a pair of wires in each muscle to record EMG activity (details below). Prior to data collection, the animal’s hindlimbs were always rigidly clamped in the stereotaxic frame with the ankle at 90° relative to the tibia, the knee at 130°, and the hip at 105°.

#### Spinal cord preparation

We transferred the animal to a stereotaxic frame (Kopf, Model 1530, Tujunga, CA, USA) where it remained until the end of experimental procedures. A dorsal laminectomy was carefully performed to expose the lumbosacral spinal cord (L4 to S2 spinal segments). The dura was incised and retracted laterally to expose the spinal cord tissue and allow for the identification of spinal segments and dorsal root entry zones. The cord was covered with warm mineral oil to prevent tissue drying.

#### Decerebration

While the deep anesthetic plane was maintained, we performed a craniotomy and made a precollicular incision with a blade. All anterior forebrain structures were removed via an aspirator and replaced with saline-soaked cotton to help blood coagulation. At this point, animals were considered to have complete lack of sentience (Silverman et al. 2005), and gaseous anesthesia was discontinued. The preparation was left to recover for about 60 minutes before data collection commenced.

### Experimental protocol

#### Force and EMG measurements

A descriptive schematic of the experimental setup is shown in Fig. 1A. Isometric force around the left ankle were measured from the TA and SOL muscles. The distal tendons of these two muscles were severed and connected with surgical sutures to linear force transducers. Care was taken to maintain the disarticulated muscles at their physiological length. The force data was low-pass filtered at 30 kHz. EMG activity was recorded from muscles around the ankle of the left hindlimb, including the TA, SOL, MG, and LG. We inserted pairs of insulated stainless steel fine wires (A-M Systems, WA, USA) in the belly of each muscle with 2 mm exposed wire at the tip. EMG data was acquired using custom-built differential amplifiers, band filtered between 0.1 Hz-30 kHz, and digitized at 10 kHz (Power-1401, CED, UK). The data was acquired through Spike2^®^ software (CED, UK) and stored on a computer for offline analysis.

#### Intracellular recordings

Single motoneurons (MNs) and interneurons were impaled and recorded in-vivo using sharp intracellular glass electrodes pulled using a micropipette puller (P97, Sutter instruments, CA). The electrode tip had a resistance of 5-15 MΩ when filled with electrolyte solution (1M K. acetate + 1 M KCl). We advanced the electrode tip into the cord tissue using a micro-stepper (Burleigh, Inchworm, model 8200). The ankle extensor and flexor motoneurons were identified by their antidromic responses to peripheral stimulation (0.1 ms pulses, Grass® stimulator, model S8800) of the tibial or common peroneal nerves, respectively. Cells were accepted for recording when they maintained a resting membrane potential below -60 mV and the antidromic spike was ≥ 60 mV. Intracellular potential was recorded using an Axoclamp-2B amplifier (Molecular Devices, CA) running in bridge or discontinuous current clamp (DCC) mode. Intracellular data was low-pass filtered at 10 kHz, digitized at 20 kHz, and acquired into Spike2 software for offline analysis.

#### Spinal cord stimulation (SCS)

Electrical stimulation of the spinal cord was generated by a custom-built voltage-controlled current source (VCCS). To ensure accurate transfer of intended stimuli, both voltage command and delivered current were simultaneously recorded at 20 kHz alongside the evoked physiologic responses. Stimulation was delivered via a custom-made bipolar silver ball electrode fitted in a spring-loaded mechanism to prevent tissue damage whenever vertical movements occurred (Fig. 1C). The electrode was placed at each location and different combinations of stimulation frequency and amplitude were repeated. The stimulation locations were set at dorsal root entry zones in different segments (Fig. 1B) including caudal end of L4 (L4C), rostral end of L5 (L5R), caudal end of L5 (L5C), rostral end of L6 (L6R), junction between L6 and L7 (L6-L7), junction between L7 and S1 (L7-S1), junction between S1 and S2 (S1-S2). In some animals, the distance between L4C and L5R or L5C and L6R was too small and was treated as a single location instead (i.e. L4-L5 junction or L5-L6 junction). The 3-D location of the identifiable dorsal root entry zones (L4-S2) as well as the border of laminectomy was recorded using a digitizer system (Microscribe G2X). These digitally-recorded locations were used to guide the placement of the stimulation electrode on the cord.

Stimulation was delivered as 1 ms monophasic square waves for 5 seconds (Fig. 1C). To determine optimum parameters needed to evoke post-stimulation activity, we used combinations of stimulation frequencies (10, 20, and 40 Hz), and amplitudes (50 - 600 µA, in increments of 50 or 100).

### Data Analysis

#### EMG data analysis

The time series data was imported from Spike2 into MATLAB and zeroed at the initiation of the 5 second stimulation train. For each channel (muscle), a window ranging from -1 to +35 seconds was extracted, the DC shift was removed, and data was rectified. For every trial, a moving average with a 300 ms window was computed from the series (movsum(), MATLAB 2023a). Then, the root mean square of the average was taken (rms(), MATLAB 2023a) from end of stimulation to end of the recording window (+5 to +35 sec) to evaluate the post-stim response.

*The effect of stimulation location* (This analysis is shown in fig. 3). To identify the optimum location, the data extracted from each experiment was grouped according to stimulation frequency, amplitude and location. The mean and standard deviation was taken for multiple trials of the same parameters (typically 3 trials for each combination). Outliers were removed as points outside of a 0-90 percentiles range (rmoutliers(), MATLAB 2023a). Experiments with less than 3 stimulation locations were excluded from this analysis. To simplify the analysis, we used 40 Hz stimulation data, and searched for maximum response to any stimulation amplitude at each location, then the stim amplitude that evoked the largest of these values was selected to be the one used across all locations. This process was repeated for each experiment separately, and the selected amplitude is displayed in the legend of fig. 3. The data was then normalized to the maximum LEAP response for each experiment. The data was split on the plot into two groups, those peak response in the lumbar segments and others which continued to show larger responses with stimulation of sacral segments.

*The effect of stimulation frequency* (This analysis is shown in fig. 4). To evaluate the effect of stimulation frequency (10, 20, 40 Hz) on LEAP response, we first extracted the data in each experiment, averaged trials with same parameters and excluded outliers as discussed above. Then the optimum stimulation location for each experiment was chosen from the analysis above to be used for comparing different frequencies. To select a stimulation amplitude, the algorithm searched for the amplitude closest to half the value of the maximum stimulation amplitude. This was done to make sure that the effect of stimulation frequency is not masked due to saturation. For each experiment, the data was normalized to the largest LEAP response at that selected stimulation amplitude.

*The effect of stimulation amplitude* (This analysis is shown in fig. 5). The data was grouped, repetitions averaged, and outliers removed using the same technique as described above. The optimum stimulation location for each experiment was chosen (from the analysis in fig. 3) to evaluate the effect of stimulation amplitude. This analysis was run separately at each frequency (10, 20, and 40 Hz). The plot in fig. 5 shows the analysis of the 40 Hz data, while analysis of the 10, and 20 Hz is in suppl. Fig. 4. The data was normalized to the maximum LEAP response for each experiment. These plots are akin to an activation curve.

#### Intracellular Data

To identify the underlying mechanism for motoneuron firing activity during LEAP, we performed intracellular recordings of ankle extensor motoneurons. After impalement, the cells were held at different potentials by injecting a DC current. The holding potential was measured as the maximum deflection in voltage following the injection of the DC current. LEAP was repeatedly evoked using the same stim parameters at each holding potential. Any evoked spiking was low-pass filtered to reveal the underlying EPSP. The peak amplitude of the evoked EPSP was measured and plotted against the holding potential. This analysis is shown in fig. 6.

## FUNDING

The study was supported by the National Institute of Health (NIH R01NS109552 and NIH R37NS135820), and Craig H. Neilsen Foundation (599050) to CJH. AM was also supported by Craig H. Neilsen Foundation fellowship (649297).

## AUTHOR CONTRIBUTIONS

CJH, AM, and MC conceived the study and designed the experiments. MC, AM, and JM designed and implemented hardware and software tools for the experiments. MJ, AM, JM, and MC performed the surgery. AM and MC performed the experiments and collected the data. AM and MC analyzed the data and generated the figures. CJH supervised the project and participated in data interpretation. The manuscript was originally written by AM and all authors contributed to its revision.

## COMPETING INTERESTS

The authors declare no competing interests.

## SUPPLMENTARY FIGURES

**Supplementary Fig. 1:**
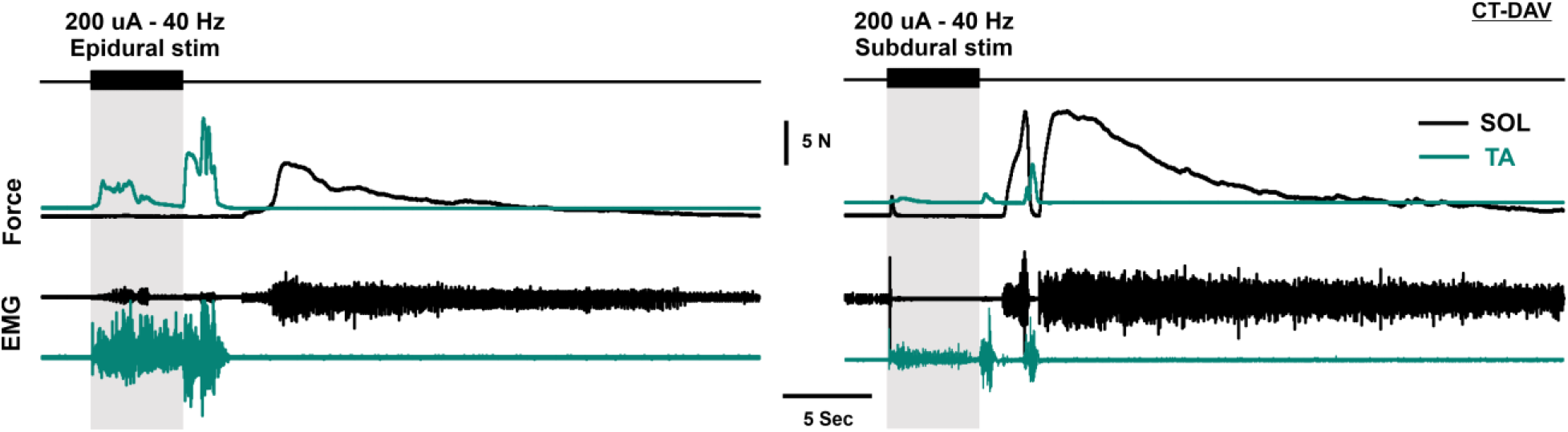
LEAP can be evoked by subdural and epidural spinal cord stimulation. Force and EMG responses of SOL and TA showing LEAP response evoked by epidural (left) and subdural stimulation (right) in the same animal. Stimulation was delivered at the same location, frequency, and amplitude before and after opening the dura, respectively (N= 5 experiments).

**Supplementary Fig. 2:**
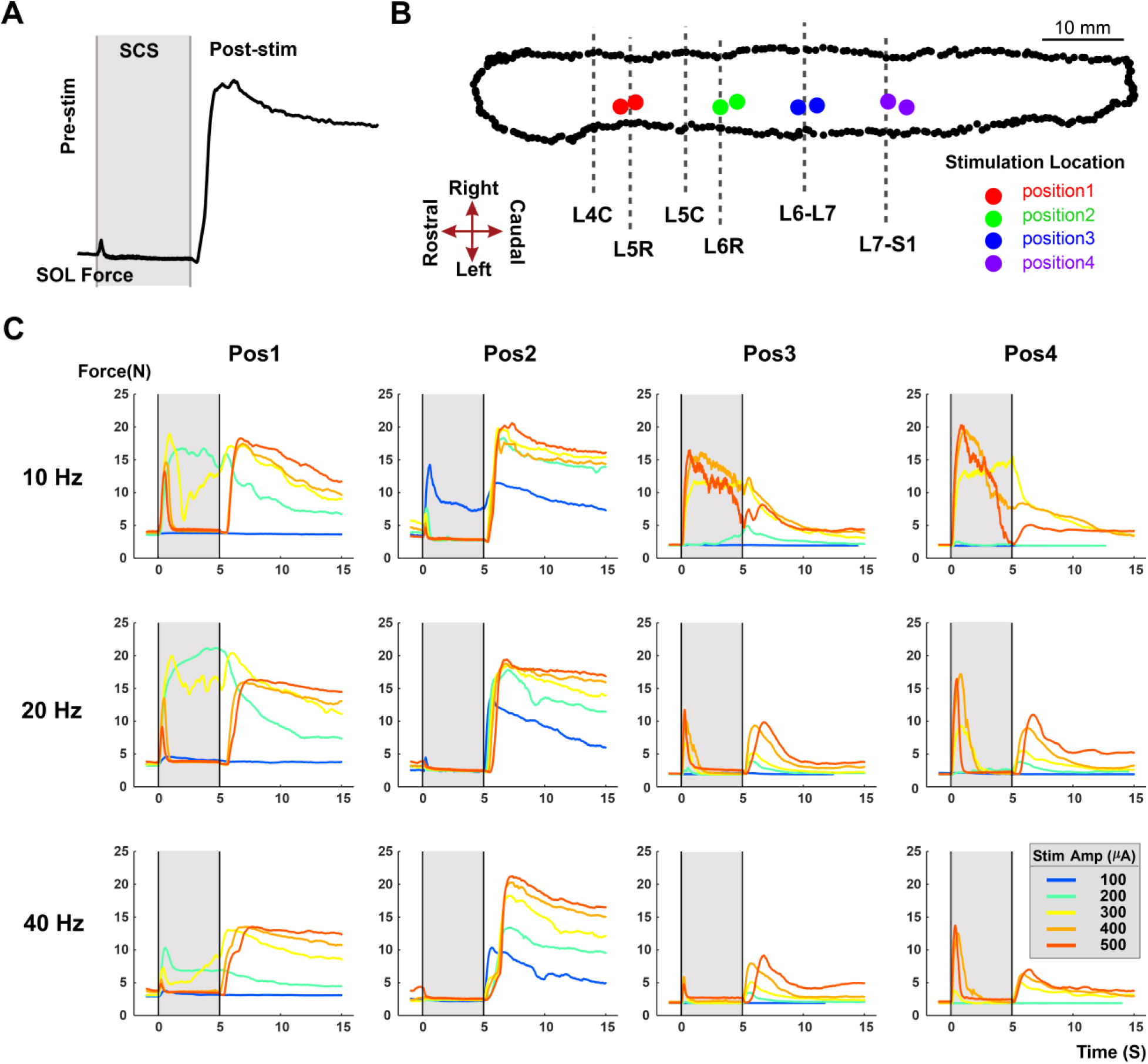
Effect of stimulation parameters on LEAP response amplitude. **A:** SOL force trace showing a typical LEAP response. Stimulation trains were delivered for 5 sec. **B:** Two-dimensional representation of the 3D digitized recording of the lumbar spinal cord surface marking dorsal root entry zones and stimulation locations recorded during the experiment using Microscribe MLX 3-D Digitizer system. The dotted black line is the border of laminectomy. **C:** Example data collected in one experiment showing multiple trials involving different combinations of stimulation positions, frequencies, and amplitudes.

**Supplementary Fig. 3.**
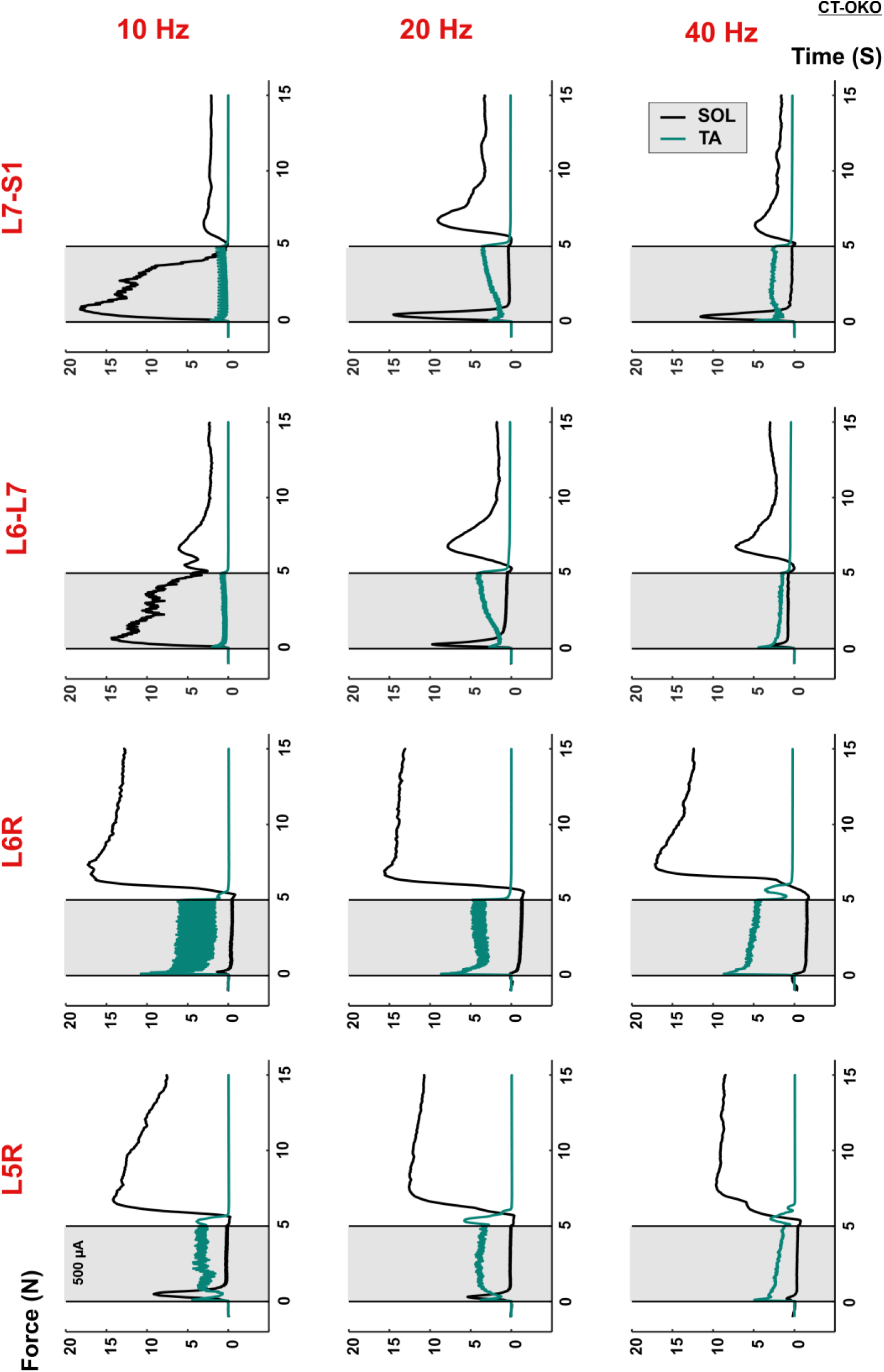
Alternating activation of SOL and TA muscles during and after stimulation. Example force traces of SOL and TA muscles from same experiment showing response to 5-sec stimulation trains at 500 µA at different locations and frequencies. Alternating activity in the antagonist muscle pair suggests the involvement of reciprocal inhibition circuits.

**Supplementary Fig. 4:**
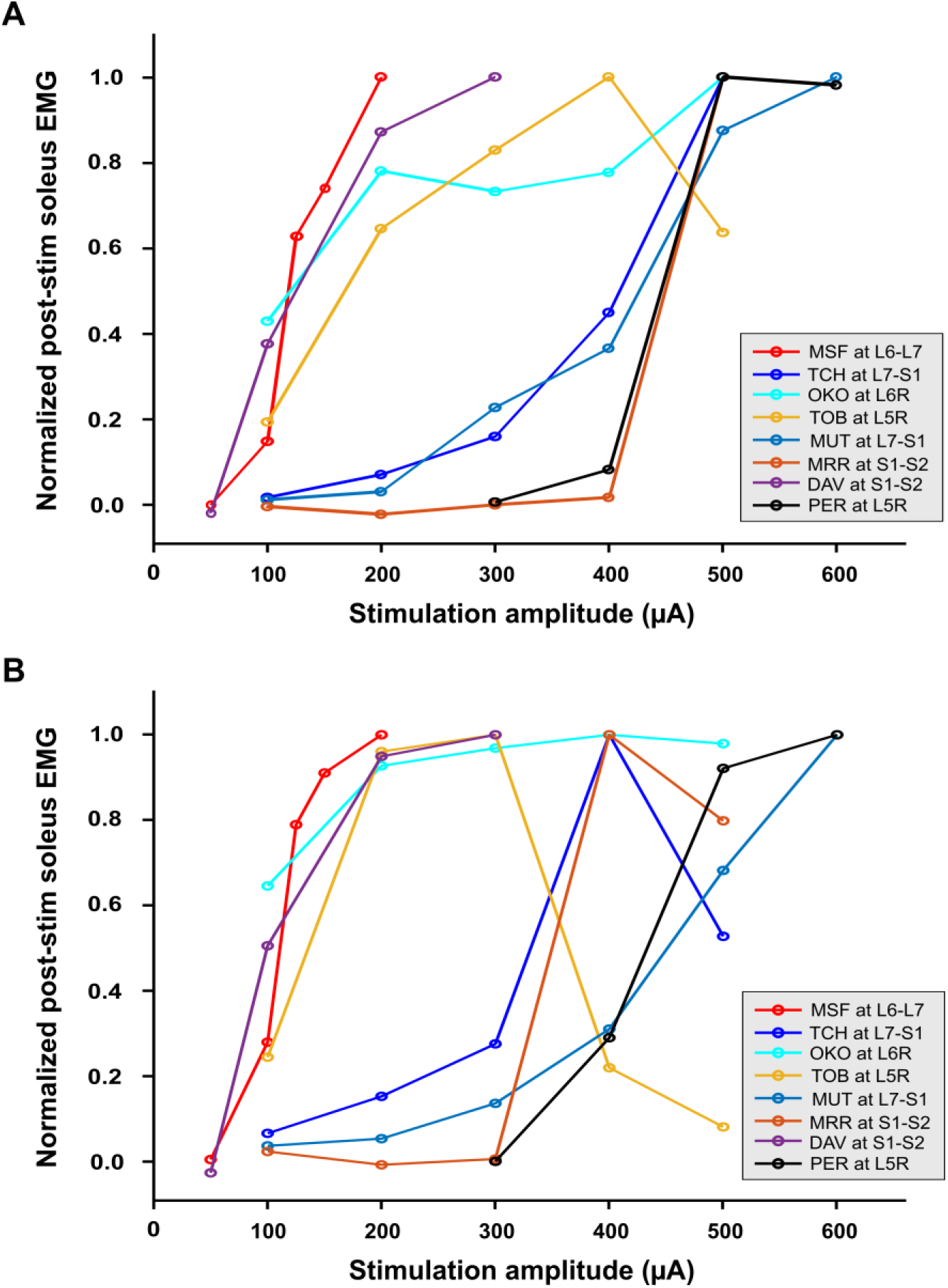
Effect of stimulation amplitude on LEAP response at different frequencies. Summary of the effect of stimulation amplitude (50-600 µA) on LEAP amplitude in 8 experiments (colors, same as in fig. 3, 4, and 5) at different frequencies. Electrical stimulation was delivered at 10 Hz (A) or 20 Hz (B) to the optimum stimulation location in each experiment (chosen from fig. 3). LEAP amplitude was measured as the integrated EMG response in soleus in the 30 seconds following the stimulation train. The data show that LEAP response can be graded using stimulation amplitude at a wide range of frequencies.

**Supplementary Fig. 5:**
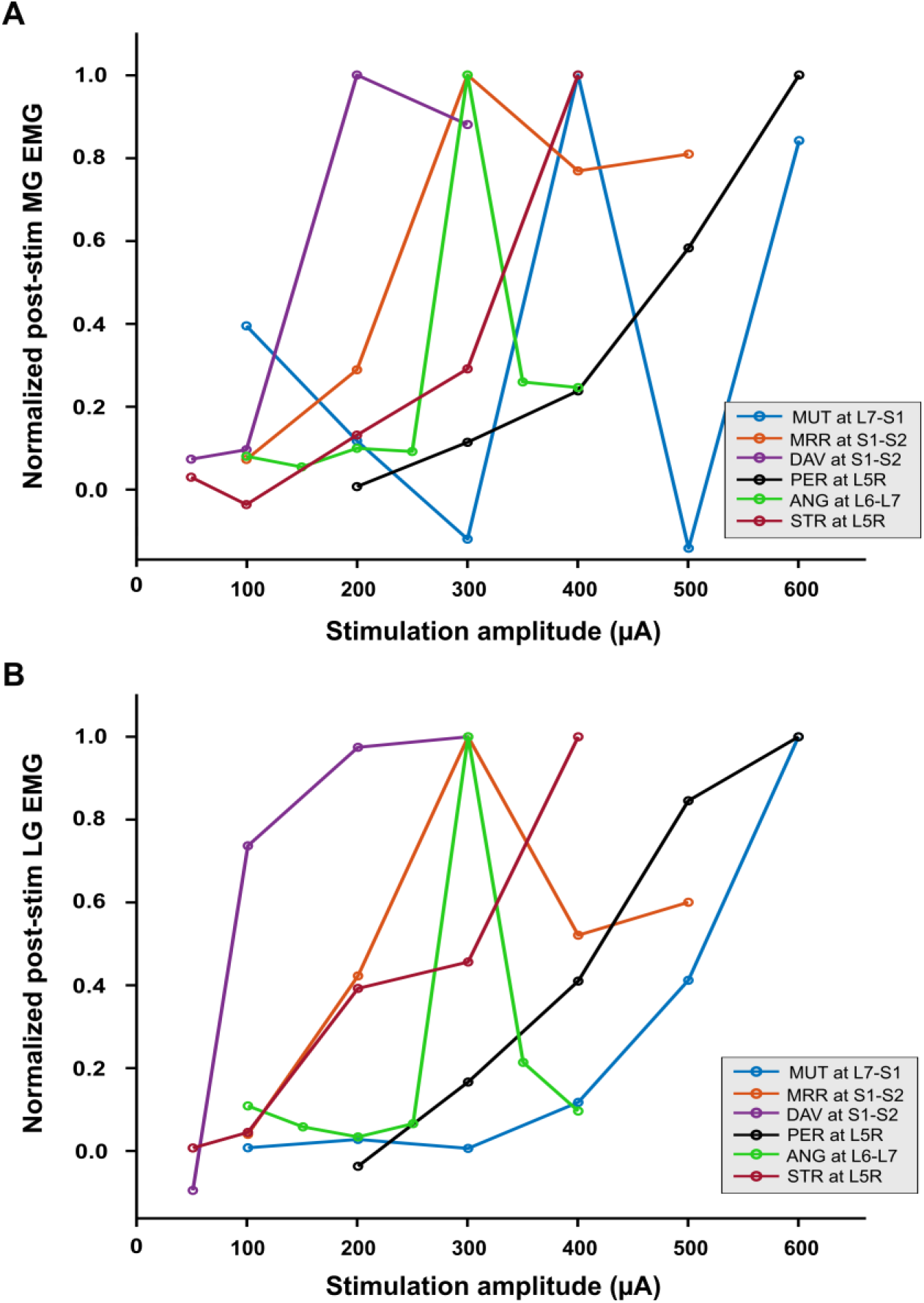
LEAP response in other ankle extensor muscles. Summary of the effect of stimulation amplitude (50-600 µA) on LEAP response in other ipsilateral triceps muscles, medial gastrocnemius (MG) and lateral gastrocnemius (LG). The plots show data from 6 experiments (same color codes as other plots). Electrical stimulation was delivered at 40 Hz to the optimum stimulation location in each experiment (chosen from fig. 3). Motor output is measured as the integrated EMG response in MG (A) or LG (B) in the 30 seconds following the stimulation train. The data show that the MG and LG muscles respond to stimulation and synergistically augment LEAP response in soleus.

## Notes

### Competing Interest Statement

The authors have declared no competing interest.

